# Fin ray branching is defined by TRAP^+^ osteolytic tubules

**DOI:** 10.1101/2022.05.09.491182

**Authors:** João Cardeira-da-Silva, Anabela Bensimon-Brito, Marco Tarasco, Ana S. Brandão, Joana Rosa, Paulo J. Almeida, António Jacinto, M. Leonor Cancela, Paulo J. Gavaia, Didier Y. R. Stainier, Vincent Laizé

**Affiliations:** Department of Developmental Genetics, Max Planck Institute for Heart and Lung Research, Bad Nauheim, Germany; DZHK German Centre for Cardiovascular Research, Partner Site Rhine-Main, Bad Nauheim, Germany; Centre of Marine Sciences (CCMAR), University of Algarve, Faro, Portugal; INSERM, ATIP-Avenir, Aix Marseille University, Marseille Medical Genetics, Marseille, France; CEDOC, NOVA Medical School, NOVA University of Lisbon, Lisbon, Portugal; STAB VIDA-Investigação e Serviços em Ciências Biológicas, Madan Parque, Caparica, Portugal; Faculty of Medicine and Biomedical Sciences (FMCB) and Algarve Biomedical Center (ABC), University of Algarve, Faro, Portugal

**Keywords:** Zebrafish, osteoclasts, osteoblasts, Sonic hedgehog, bone patterning, bone resorption, bone formation, bifurcation, fin regeneration, fin development

## Abstract

The shaping of bone structures relies on various cell types and signalling pathways. Here, we use the zebrafish bifurcating fin rays during regeneration to investigate bone patterning. We found that the regenerating fin rays form via two mineralization fronts that undergo an osteoblast-dependent fusion/stitching until the branchpoint, and that bifurcation is not simply the splitting of one unit into two. We identified tartrate-resistant acid phosphatase-positive (TRAP^+^) osteolytic tubular structures at the branchpoints, here named osteolytic tubules (OLTs). Chemical inhibition of their bone-resorbing activity strongly impairs ray bifurcation, indicating that OLTs counteract the stitching process. Finally, by testing different osteoactive compounds, we show that the position of the branchpoint depends on the balance between bone mineralization and resorption activities. Overall, these findings provide a new perspective on fin ray formation and bifurcation, and reveal a key role for OLTs in defining the proximo-distal position of the branchpoint.

**Graphical summary:** 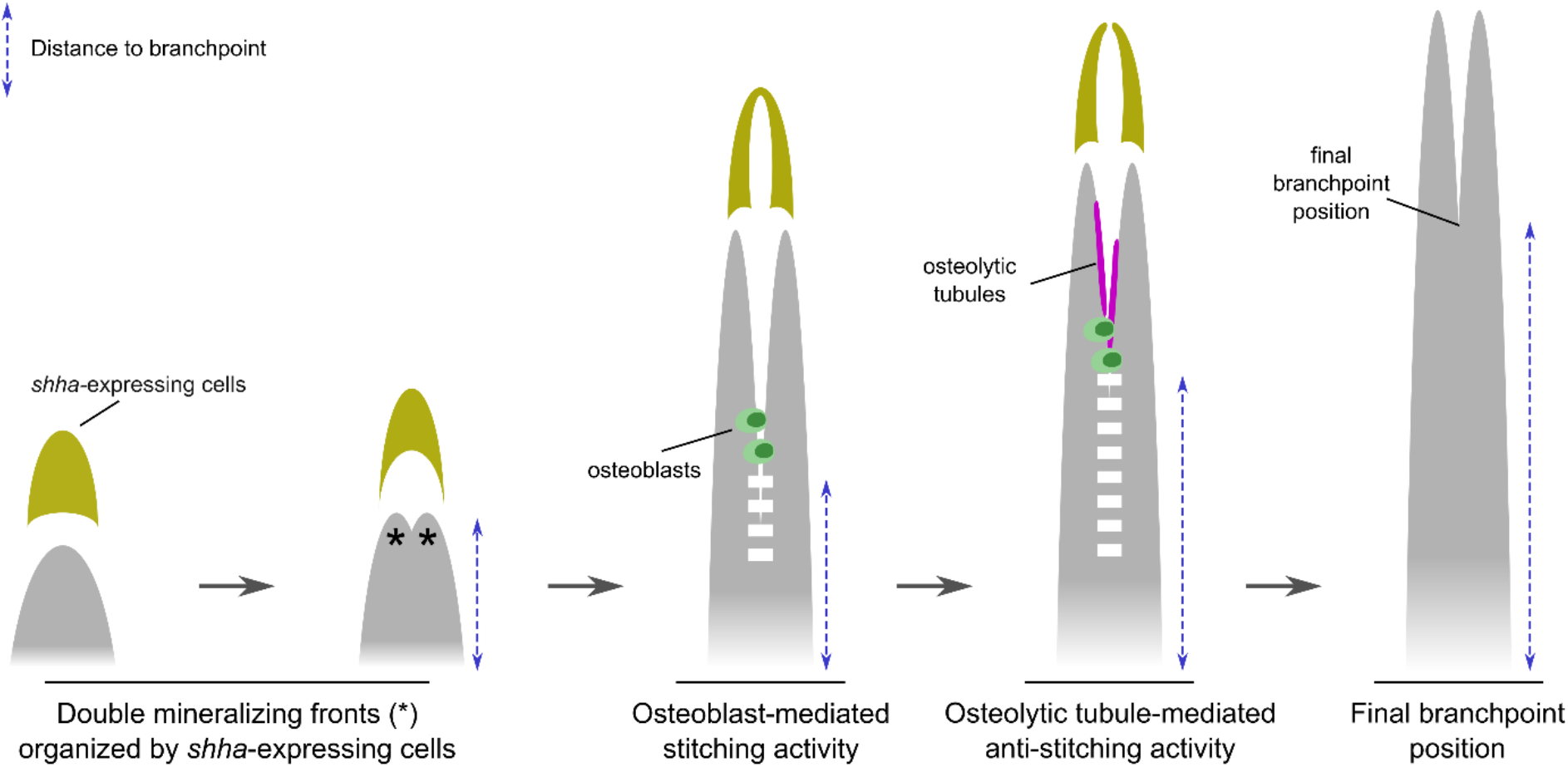

## INTRODUCTION

The formation of skeletal structures, such as the limbs or the vertebral column, entails precise patterning processes orchestrated through the activity of multiple cells and signaling pathways (Salhotra et al., 2020). Bone formation and homeostasis depend on the synchronized activity of bone-forming (osteoblasts) and -resorbing (osteoclasts) cells (Kim et al., 2020). Understanding how these cells contribute to the shaping of bones is particularly relevant in the context of bone disease and injury (Salhotra et al., 2020).

Although mammals have a limited capacity to regenerate skeletal tissues, except for the digit tips (Fernando et al., 2011), non-mammalian vertebrates, such as zebrafish (*Danio rerio*), are able to regenerate multiple organs and appendages very efficiently (Gemberling et al., 2013). In this regard, zebrafish is an established model for studying caudal fin regeneration, during which multiple tissues, including bone, are fully restored to their original size and shape (Akimenko et al., 2003; Gemberling et al., 2013; Pfefferli and Jaźwińska, 2015; Sehring and Weidinger, 2020). The process of fin regeneration involves the formation of a blastema, a pool of dedifferentiated cells that form the new tissues, distal to each ray (Pfefferli and Jaźwińska, 2015; Stewart and Stankunas, 2012). The fin bones, or fin rays (lepidotrichia), are composed of a mineralized collagenous matrix secreted by osteoblasts and are formed by dermal ossification without a cartilaginous template (Becerra et al., 1983; Marí-Beffa et al., 1996; Montes et al., 1982). Fin rays are composed of repetitive segments in the proximo-distal axis and by two hemirays (one left and one right) in the transverse plane. Apart from the outermost ones, all rays are bifurcated at different proximo-distal positions, according to the shape of the fin and ray length, i.e., shorter rays branching more proximally and longer rays more distally (Cardeira et al., 2016). The proximo-distal positioning of the bifurcations along the rays is critical for the overall fin architecture, as recently highlighted by studies of *collagen9ac1c* zebrafish mutants which display impaired bifurcation associated with fin misshaping (Nakagawa et al., 2022).

Ray bifurcation is considered as the splitting of a single mineralizing ray into two daughter rays forming a branched structure. This process depends on various signals including those from the adjacent interray tissue (Murciano et al., 2001, 2002). Importantly, among the signaling pathways controling bone patterning, such as during limb development (Capdevila and Belmonte, 2001; Riddle et al., 1993), Sonic hedgehog (Shh) has been described as a key factor controling ray bifurcation during fin regeneration (Armstrong et al., 2017; Laforest et al., 1998). However, repetitive amputations (Azevedo et al., 2012), macrophage activity (Petrie et al., 2014), actinotrichia organization (Nakagawa et al., 2022), and hydrodynamics (Dagenais et al., 2021) are able to alter the proximo-distal position of ray branchpoints through apparently Shh-independent mechanisms. Thus, the formation and shaping of bifurcated bony structures and the cellular contribution to these processes remain unclear.

Particularly, the structural events of fin ray formation and bifurcation, and how different cells coordinate to shape fin rays, are still unknown.

Here, we used chemical and genetic tools, together with live-imaging approaches, to investigate how caudal fin rays mineralize and are shaped during regeneration. We show that fin rays mineralize in two lateral fronts that undergo an osteoblast-dependent fusion (stitching) in the center. This stitching process is then counteracted by the activity of tartrate-resistant acid phosphatase-positive (TRAP^+^) osteolytic tubules (OLTs), which define the proximo-distal position of the branchpoints through localized osteolytic activity. Overall, we i) provide a detailed analysis of how bony rays are formed and bifurcate, ii) identify OLTs as critical in defining the branchpoint position, and iii) propose this model as instrumental to study the (im)balance between bone mineralizing and resorbing activites.

## RESULTS

### Rays form via two mineralizing fronts that undergo gradual stitching

To understand the morphogenetic processes underlying ray formation and bifurcation, we tracked the mineralization of individual rays over time in the adult zebrafish caudal fin post-amputation. To ensure consistency, we define “bifurcation” as the process of mineralized ray splitting, and “bifurcating ray” as the structure characterized by two distinguishable daughter rays forming a branchpoint (Figure S1). We found that the onset of bifurcation is variable amongst individuals, taking place between 3 and 6 days post-amputation (dpa; Figure S2). Therefore, in the present study, we distinguish the “pre-bifurcation” and “bifurcation” stages irrespective of the post-amputation time point. We observed that, during the bifurcation phase, the branchpoints are positioned at more proximal sites (i.e., closer to the amputation plane) in early stages, but gradually distalize over time (i.e., move further away from the amputation plane; Figures 1A-1B). These findings indicate that the bony rays form through two mineralizing fronts that undergo a proximal-to-distal fusion, a process herein termed “stitching”. The stitching of bifurcating rays is clear until at least 7 dpa, when it gradually slows down and the branchpoint position stabilizes at around 9-10 dpa (Figure 1B and Table S1). Moreover, we frequently observed longitudinal gaps in the mineralized rays proximal to the branchpoint, possibly corresponding to remnant imperfections of the stitching process (Figure 1C).

**Figure 1.**
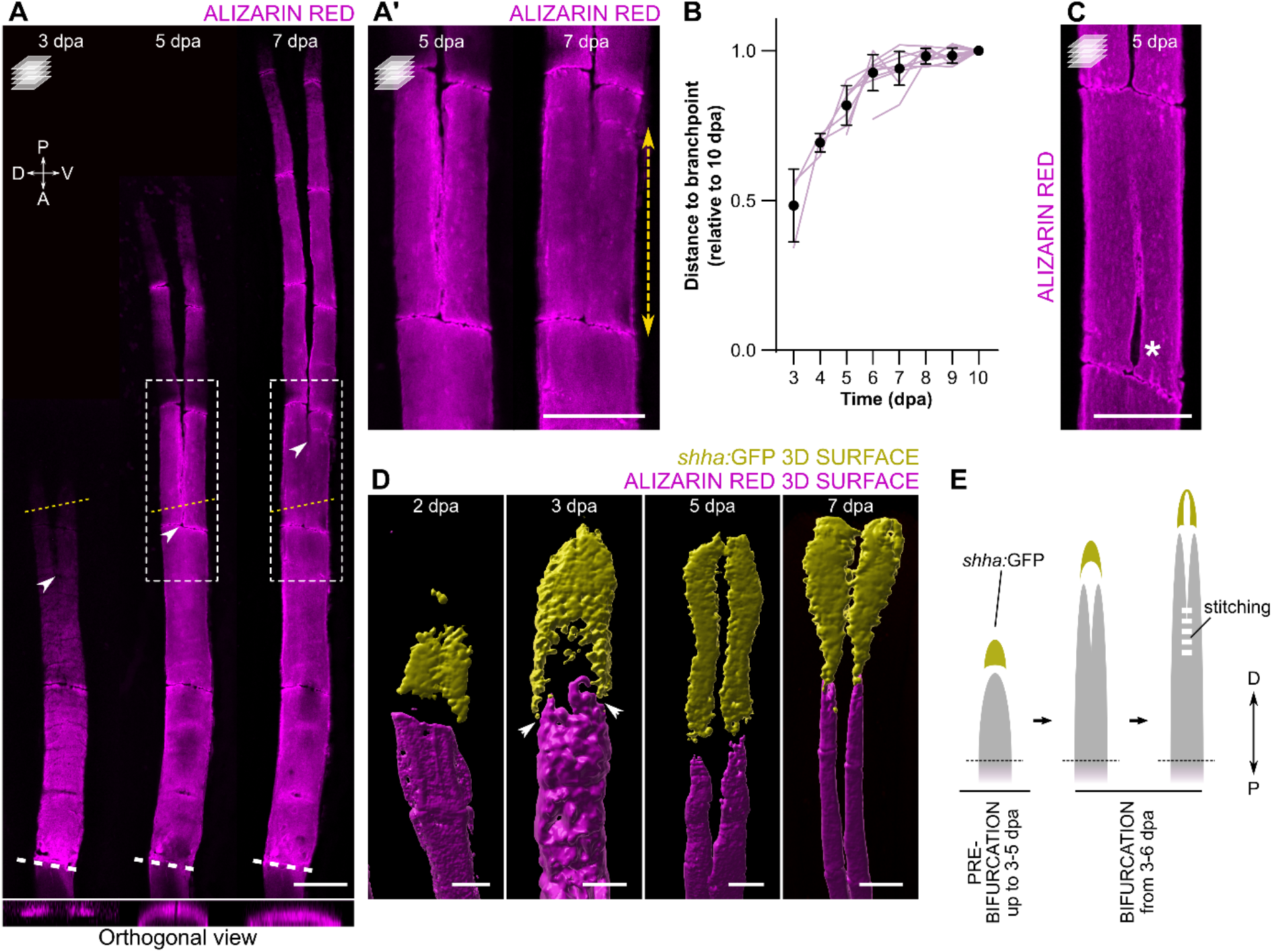
During bifurcation, rays form via two mineralizing fronts that undergo gradual stitching. (**A, A’**) Confocal images of alizarin red-stained fish, showing the regenerative progression of a single fin ray and the dynamic position of the branchpoint (arrowheads) over time. Yellow dotted lines indicate the planes of the orthogonal views on the bottom; white dashed lines indicate the amputation plane; boxes represent the magnified areas in **A’**. Double arrow shows the extension of mineral stitching from 5 to 7 dpa. (**B**) Individual quantitative analysis of the distance from the amputation plane to the branchpoint of dorsal ray #3 at different time points relative to 10 dpa (tracking of individual rays). *N* = 3 for 3 dpa, 4 for 4 dpa, 7 for 5 dpa and 10 for the remaining time points. One-way ANOVA with Geisser-Greenhouse correction (*P* < 0.0001) and Tukey post-hoc tests for multiple comparisons (details of statistics for multiple comparisons can be found in Table S1). The graph shows the mean ± SD and values of individual rays (magenta lines). (**C**) Confocal image of bifurcating ray displaying an unstitched portion (asterisk) below the branchpoint. (**D**) 3D surface renderings of regenerating rays at different time points, showing the segregation of *shha:*GFP^+^ domains proximal to the forming bone, even before the onset of bifurcation. (**E**) Model of ray regeneration divided into the pre-bifurcation and bifurcation phases, highlighting the timing variability of the different events between specimens, and showing the process of stitching of the two mineralizing fronts. P, proximal; D, distal. All confocal images are maximum intensity projections. Scale bars: 100 μm.

We then analyzed *sonic hedgehog a* (*shha*) expression with single-cell resolution imaging and observed that prior to the formation of the two mineralizing fronts, *shha:*GFP^*+*^ cells organized in a single cluster with a cap-like shape (Figure 1D). Interestingly, we observed that the overhangs of this structure were on top of the two mineralizing fronts. These data suggest that the mechanisms of ray segregation are active before the two mineralizing fronts are visible, and that regenerating rays have a predisposition to bifurcate. With the progression of the regenerative and bifurcation processes, *shha*-expressing cell domains become further extended proximal to the mineralizing fronts while remaining as a single domain until complete separation takes place around 7 dpa (Figure 1D), when the stitching process slows down towards its conclusion.

Overall, we show that, at least during part of the outgrowth, bony rays regenerate via two mineralizing fronts that progress in tandem and undergo a proximal-to-distal stitching up to the final position of the branchpoint (Figure 1E).

### Osteoblasts mediate the stitching of the two mineralizing fronts

To better understand the mechanisms and biological processes taking place during ray regeneration and bifurcation, we performed RNA-Seq on regenerates at 1 dpa (i.e., prior to new bone mineralization; Cardeira et al., 2016) and 3 dpa (i.e., at, or shortly preceding, the onset of two mineralizing fronts and cap-like *shha* single domain; Figure S3A). As expected, while various early blastema marker genes (e.g., *msx2b, fgf20a, wnt5a*) were downregulated, several osteogenesis- and morphogenesis-related genes (e.g., *sp7, col10a1a, bglap, alpl, hoxb13a, hoxd13a*) were upregulated at 3 dpa in comparison with 1 dpa (Figure S3B). Moreover, gene ontology terms related to extracellular matrix (ECM) were shown to be enriched at 3 dpa (Figure S3C). These ECM proteins include collagens and other structural components, as well as ECM regulators (Figure S3D). Importantly, osteoblast-related genes, such as *bone gla protein* (*bglap*) and *sp7 transcription factor* (*sp7*), and genes encoding bone matrix components, namely *collagen 1* (e.g., *col1a1a, col1a1b, col1a2*) and *collagen 10* (e.g., *col10a1a*), were upregulated at 3 dpa (Figure S3E).

With these data, we hypothesized that osteoblasts, known to drive the formation of new bone in regenerating fin rays (Knopf et al., 2011; Singh et al., 2012), are also involved in bone patterning and mediate the stitching process. Accordingly, by using *bglap:GFP* reporter fish, we observed osteoblasts at the stitching sites (Figure 2A). To determine the role of osteoblasts during the stitching process, we performed cell-specific ablation using the *sp7:*mCherry-NTR line (Singh et al., 2012). Zebrafish were exposed to Mtz for 24 hours, starting at 3 dpa, to maximize the depletion of osteoblasts (Figure 2B). At 7 dpa, Mtz-treated fish displayed proximalized branchpoints when compared with control fish (Figure 2C and 2D). These results indicate an impaired stitching of the two mineralizing fronts of each ray and confirm the importance of osteoblasts in mediating this process (Figure 2E).

**Figure 2.**
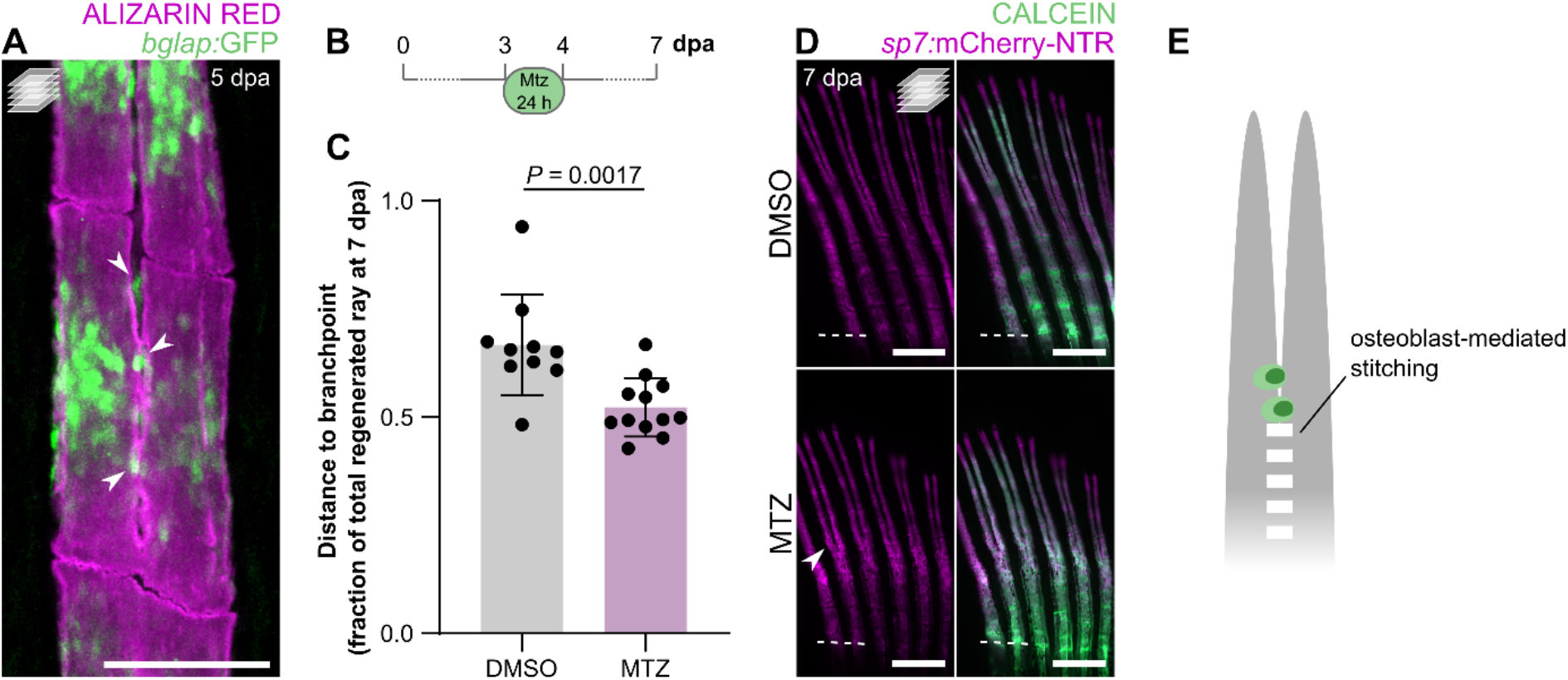
Osteoblast ablation induces proximalization of branchpoints. (**A**) Confocal image of a bifurcating alizarin red-stained ray, showing mature osteoblasts at the stitching zone (arrowheads). (**B-D**) Ablation of *sp7:*mCherry-NTR^+^ osteoblasts during early bifurcation. (**B**) Ablation protocol. (**C**) Quantification of the relative distance from the amputation plane to the branchpoint at 7 dpa, following Mtz treatment. The graph dots represent individual fish, and the bars represent the mean ± SD. Unpaired, two-tailed Student’s t-test. (**D**) Confocal images of regenerates of Mtz-treated compared with control/DMSO-treated fish, showing proximalized branchpoints (arrowhead). (**E**) Model placing osteoblasts as key players in the stitching process. Dashed white lines in **D** mark the amputation planes. All confocal images are maximum intensity projections. Scale bars: 100 μm in **A**; 500 μm in **D**.

### Osteolytic tubules are central to ray bifurcation and outgrowth

Osteoblasts and osteoclasts display opposing, yet cross-balanced, activities in numerous settings of bone formation and homeostasis (Salhotra et al., 2020). In fact, our RNA-Seq data show that together with osteoblast-, bone formation- and bone matrix-related genes, regulators of osteoclastogenesis, such as *tumor necrosis factor receptor superfamily, member 11a, NFKB activator* (*tnfrsf11a*) and *colony stimulating factor 1 receptor b* (*csf1rb*), were also upregulated at 3 dpa (Figure S3E). Therefore, we analyzed the osteoclast dynamics during the bifurcation process. By using the *ctsk:DsRed* line, which labels active mature osteoclasts (Caetano-Lopes et al., 2020), we found that *cstk:*DsRed^*+*^ cells display distinct morphologies and localizations at the different phases of ray bifurcation (Figure 3). In the pre-bifurcation phase, during the stitching process, osteoclasts are round, exhibit cytoplasmic extensions, and accumulate mostly inside the regenerating rays (i.e., in between the two hemirays; Figure 3A and 3B). In contrast, during the bifurcation phase, osteoclasts form elongated structures that line with the branching ray surface and border the outer limits of the regenerating bone (Figure 3C and 3D). To confirm that these elongated structures display bone resorbing activity, we stained the regenerating fins at the bifurcation phase for tartrate-resistant acid phosphatase (TRAP) activity, an osteoclast-specific secreted enzyme involved in bone resorption (Witten and Villwock, 1997). We observed that TRAP signal strongly accumulates at the branching sites (Figure 3E) and co-localizes with the *ctsk:*DsRed^+^ cells (Figure 3F), confirming that these cells indeed display bone-resorbing activity. We also observed that these *ctsk:*DsRed^+^ elongated structures exhibit a non-fluorescent lumen and, therefore, we name them osteolytic tubules (OLTs). Interestingly, TRAP signal strongly accumulates within the lumen of OLTs (Figure 3F).

**Figure 3.**
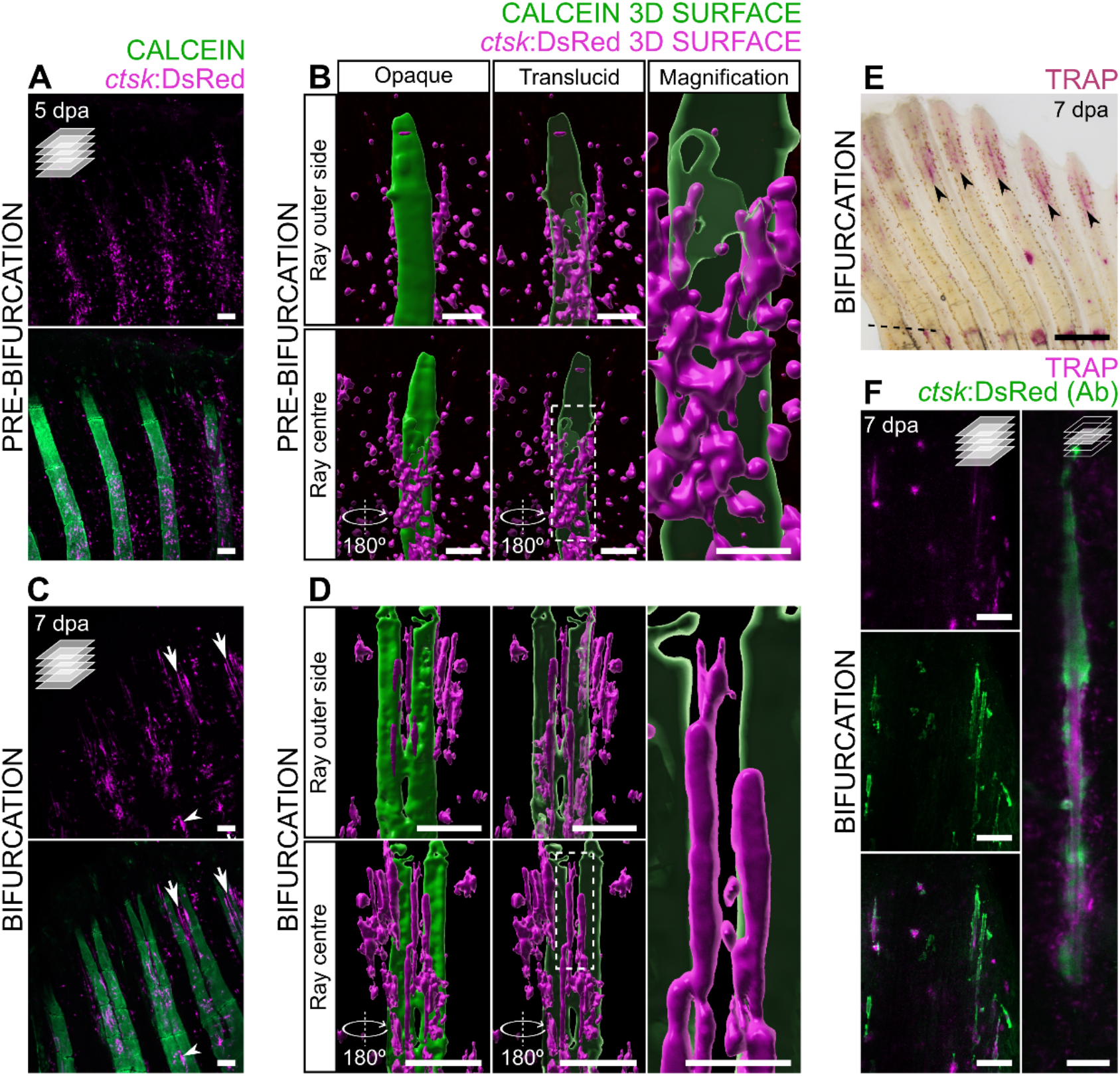
TRAP-secreting osteolytic tubules align with the bone surface at ray splitting regions during bifurcation. (**A-D**) Confocal images (**A, C**) and 3D surface renderings (**B, D**) of *ctsk*:*DsRed* expression in calcein-stained regenerating fins at the pre-bifurcation (**A, B**) and bifurcation (**C, D**) stages. 3D surface renderings are shown in two different perspectives and mineralized surface is shown as opaque and translucid; 3D reconstructions represent one hemiray and the centre of ray, and not the entire ray. Boxes represent the magnified areas on the right. (**E**) Stereomicroscope image showing TRAP activity at the splitting regions (arrowheads) of bifurcating rays. Dashed line marks the amputation plane. (**F**) Confocal images of elongated osteoclasts at the splitting regions, immunostained for DsRed and stained for TRAP activity. Single confocal planes in F right; all remaining confocal images are maximum intensity projections. Scale bars: 100 μm in **A-D** (lower magnifications in **B** and **D**); 30 μm in higher magnifications in **B** and **D**; 500 μm in **E**; 50 and 10 μm in **F** left and right, respectively.

To understand whether OLTs are specific to regeneration, we analyzed uninjured fins, which display little *ctsk:*DsRed signal, with a few OLTs at the distal tips of the permanently growing fin rays (Figure 4A). This observation indicates that OLTs are not specific to the regenerative process and are associated with ray formation in general. In fact, OLTs are also abundant in developing fins (Figure 4B and 4C). To further characterize the OLTs, we assessed the spatial co-localization of the expression of different reporter transgenes in developing and regenerating fins. Because osteoclasts and macrophages share a common lineage, we imaged OLTs in the *mpeg1*.*1:YFP* background, and observed that they form distinct populations. However, they seem to physically interact (Figure 4D), suggesting that the immune system may play a role in regulating the OLTs and/or vice-versa. Using the *fli1:GFP* reporter line, we observed that OLTs align in close proximity to blood vessels, both during development and regeneration (Figure 4E), also suggesting a relationship between OLTs and the vasculature. Interestingly, we observed that OLTs express the *lyve1b:GFP* lymphatic endothelial reporter (Figure 4F).

**Figure 4.**
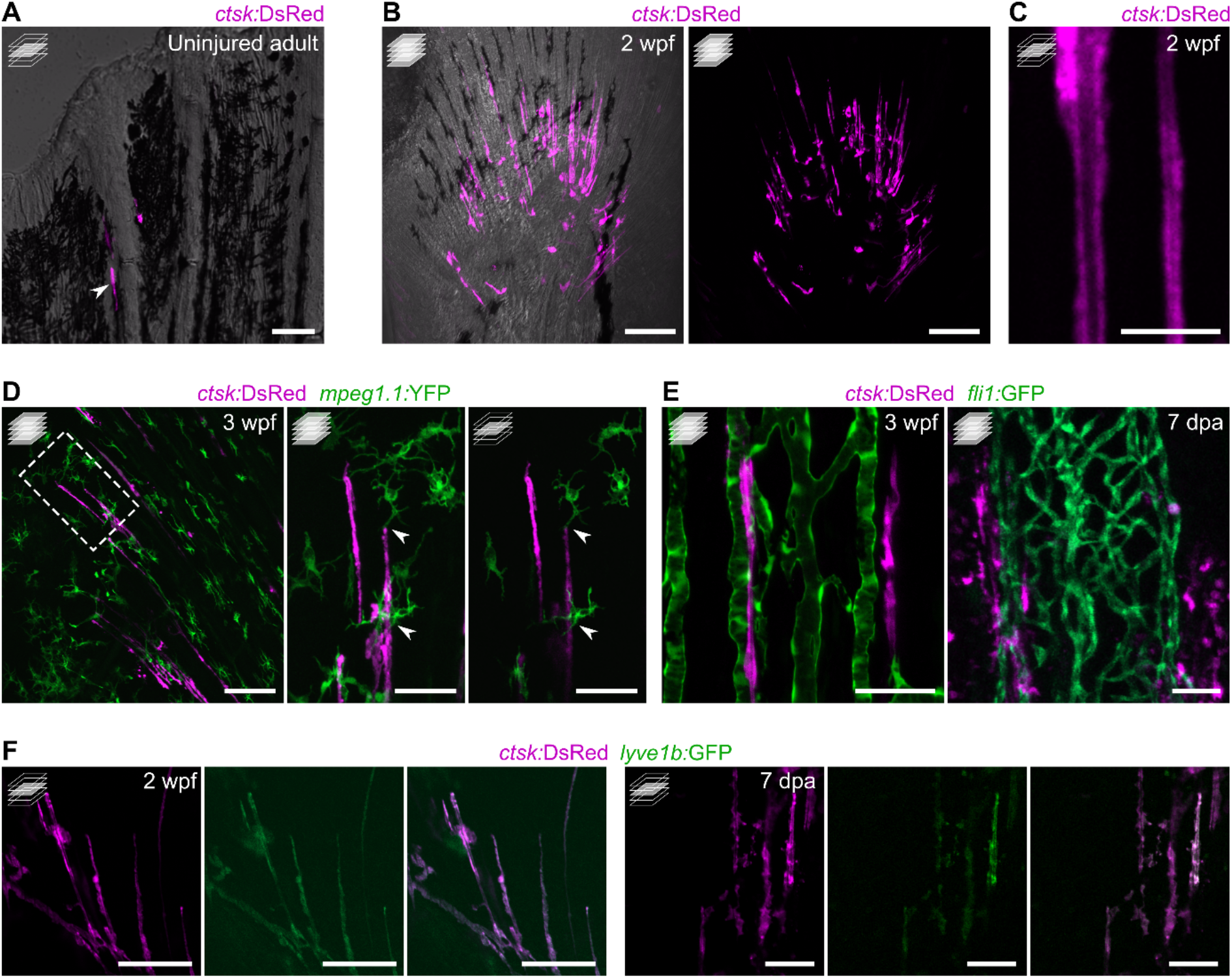
Osteolytic tubules are present in the developing fin, display a tubular morphology, interact with macrophages, align with blood vessels and are positive for *lyve1b*:*GFP*. (**A**) Confocal image of adult uninjured caudal fin displaying *ctsk*:DsRed^+^ osteoclasts at the growing edges of rays. (**B, C**) Confocal images of developing caudal fin showing elongated osteoclast structures aligned with fin rays (**B**) displaying a tubular morphology (**C**). (**D**) Confocal images of developing pectoral fin, showing direct interaction between macrophages (green) and tubular osteoclasts. Box represents the magnified areas on the right. (**E**) Confocal images showing tubular osteoclasts aligning with blood vessels (green) during caudal fin development and regeneration (left and right, respectively). (**F**) Confocal images showing that tubular osteoclasts are positive for *lyve1b*:*GFP*, during the development and regeneration (sets of images on the left and right, respectively) of the caudal fin. Maximum intensity projections in B, D (lower magnification and higher magnification on the left) and E. Single confocal planes in A, C, D (higher magnification on the right) and F. Scale bars: 50 μm in A, D (higher magnifications), E and F; 100 μm in B and D (lower magnification); 10 μm in C.

These data show the presence of OLTs during bone development and regeneration, particularly in association with the bifurcation process of regenerating rays, possibly interacting with the immune and vascular systems.

### Bone resorbing activity of OLTs is required for bifurcation

Given the presence of OLTs at the branching sites, we investigated whether they were required during the ray bifurcation process. A single dose of salmon calcitonin, a strong anti-resorbing agent (Chesnut et al., 2008), also active in fish (Mukherjee et al., 2004; Suzuki et al., 2000), was injected intraperitoneally simultaneously to the amputation procedure, and ray regeneration was tracked over time (Figure 5A). We observed a strong impairment of ray bifurcation in calcitonin-injected fish, as opposed to control fish injected with DMSO (Figure 5B), as shown by a distalization of the branchpoints (Figure 5C) and a reduction in the number of rays displaying signs of bifurcation at 7 dpa (Figure S4A), in all concentrations tested. At 14 dpa, when the regenerative process is close to conclusion, the branchpoint distalization was still markedly evident (Figure S4B), with some fish presenting no bifurcated rays at all. Because the highest calcitonin concentration (50 μg.g^-1^) showed high phenotypic variability and heterogeneities in mineralization density (i.e., variable ARS staining intensities within fins; Figure S4C), and the lowest concentration (0.5 μg.g^-1^) was enough to induce a bifurcation phenotype, subsequent analyses were carried out at 0.5 μg.g^-1^ to avoid unspecific effects of the drug. To show that the calcitonin treatment was indeed inhibiting bone resorption activity, we analyzed TRAP accumulation at 5 dpa and observed a significant reduction in the treated fish (Figure 5D). The number of rays displaying TRAP signal in the bifurcation site was also drastically reduced in calcitonin-treated zebrafish (Figure 5E), confirming the effectiveness of this drug in reducing bone resorbing activity. Interestingly, inhibition of bifurcation and distalization of the branchpoints were not accompanied by an impaired segregation or length of the *shha:*GFP^+^ domains (Figure 5F, G). These results indicate that, apart from the role of Shh signaling in mobilizing pre-osteoblasts and inducing ray splitting (Armstrong et al., 2017), the positioning of branchpoints is not dependent on Shh signaling but requires additional cellular and molecular players.

**Figure 5.**
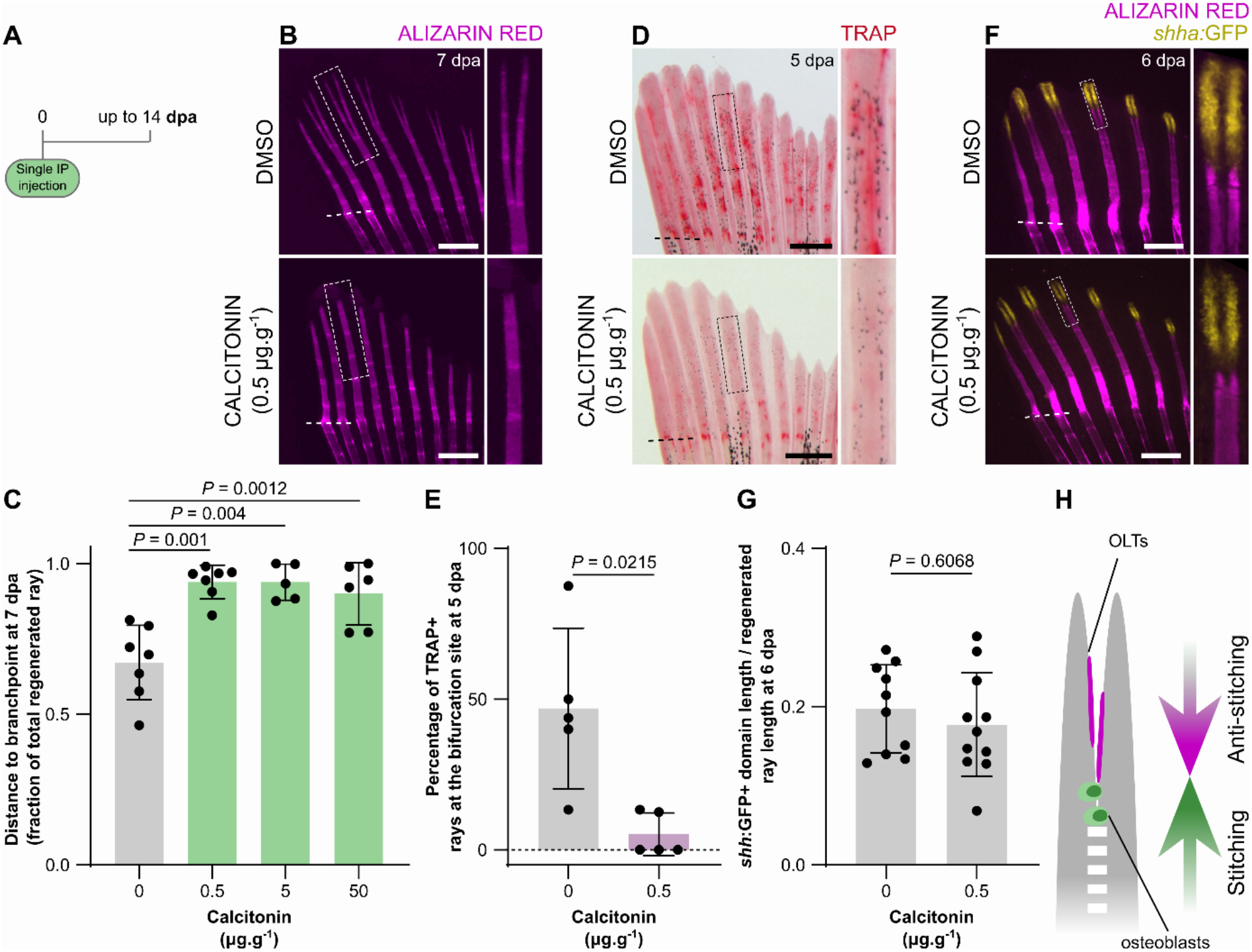
Inhibition of osteoclast activity impairs ray bifurcation. (**A**) Calcitonin treatment experimental protocol. (**B, C**) Stereomicroscope images of mineralized rays (B) and quantification (C) of the relative distance from the amputation plane to the branchpoint at 7 dpa, showing impaired bifurcation upon calcitonin treatment. Boxes in B represent the magnified areas. (**D, E**) Stereomicroscope images of TRAP-stained fins (D) and quantification of the percentage of rays displaying TRAP signal at the bifurcation site (E) at 5 dpa, showing reduced osteoclast activity upon calcitonin treatment. Boxes in D represent the magnified areas. (**F, G**) Confocal images of bifurcating rays showing segregating *shha:*GFP^*+*^ domains (F) and quantification of their extension (G), showing that calcitonin does not affect Shh signaling. Boxes in F represent the magnified areas. (**H**) Model highlighting the balanced stitching and anti-stitching activities of osteoblasts and OLTs to define the branchpoint position. All graphs show the mean ± SD. One-way ANOVA (*P* = 0.0057) and Tukey’s post-hoc test in C; unpaired, two-tailed Student’s t-test with Welch’s correction in E; unpaired, two-tailed Student’s t-tests in F. Scale bars: 500 μm.

Taken together, our data show that bone resorbing activity of OLTs is central to ray bifurcation, and we propose that OLTs are responsible for the anti-stitching activity to counteract osteoblast-mediated stitching and, thus, define the branchpoints of forming rays (Figure 5H).

### Branchpoints as indicators of bone resorption/mineralization imbalances

After establishing the cooperative action of osteoblasts and OLTs in defining the branchpoints, we evaluated whether the bifurcation process could represent an adequate readout of imbalanced activities of these cell types. For that end, we exposed amputated fish to different drugs with known pro-resorbing (dexamethasone and prednisolone; Hardy et al., 2018), pro-mineralogenic (ibandronate; Bauss and Dempster, 2007) or osteotoxic properties when in excess (retinoic acid; Conaway et al., 2013). The corticosteroids dexamethasone and prednisolone were shown to induce a proximalization of the branchpoints at 5 dpa (Figures 6A-6C) indicating decreased stitching and/or increased anti-stitching activity. The highest concentration of prednisolone even induced trifurcations (Figure 6A), suggesting that the bone rays may have more than one simultaneous branchpoint leading to skeletal malformations. In parallel, we tested the effects of retinoic acid, known to disrupt osteoblast boundaries and to block osteoclast differentiation during fin regeneration (Blum and Begemann, 2015a, 2015b). As observed with calcitonin, retinoic acid promoted a distalization of the branchpoints and induced an almost complete blockage of bifurcation still at 7 dpa (Figures 6D and 6E), likely by increased stitching and/or decreased anti-stitching activity. However, this drug also results in deformed and shorter regenerating rays, which may cause a delay in bifurcation. Therefore, we tested ibandronate, a bisphosphonate widely used as anti-resorbing drug, which also resulted in distalized branchpoints, as seen at 5 dpa (Figures 6F and 6G). This result indicates increased stitching and/or decreased anti-stitching activity.

**Figure 6.**
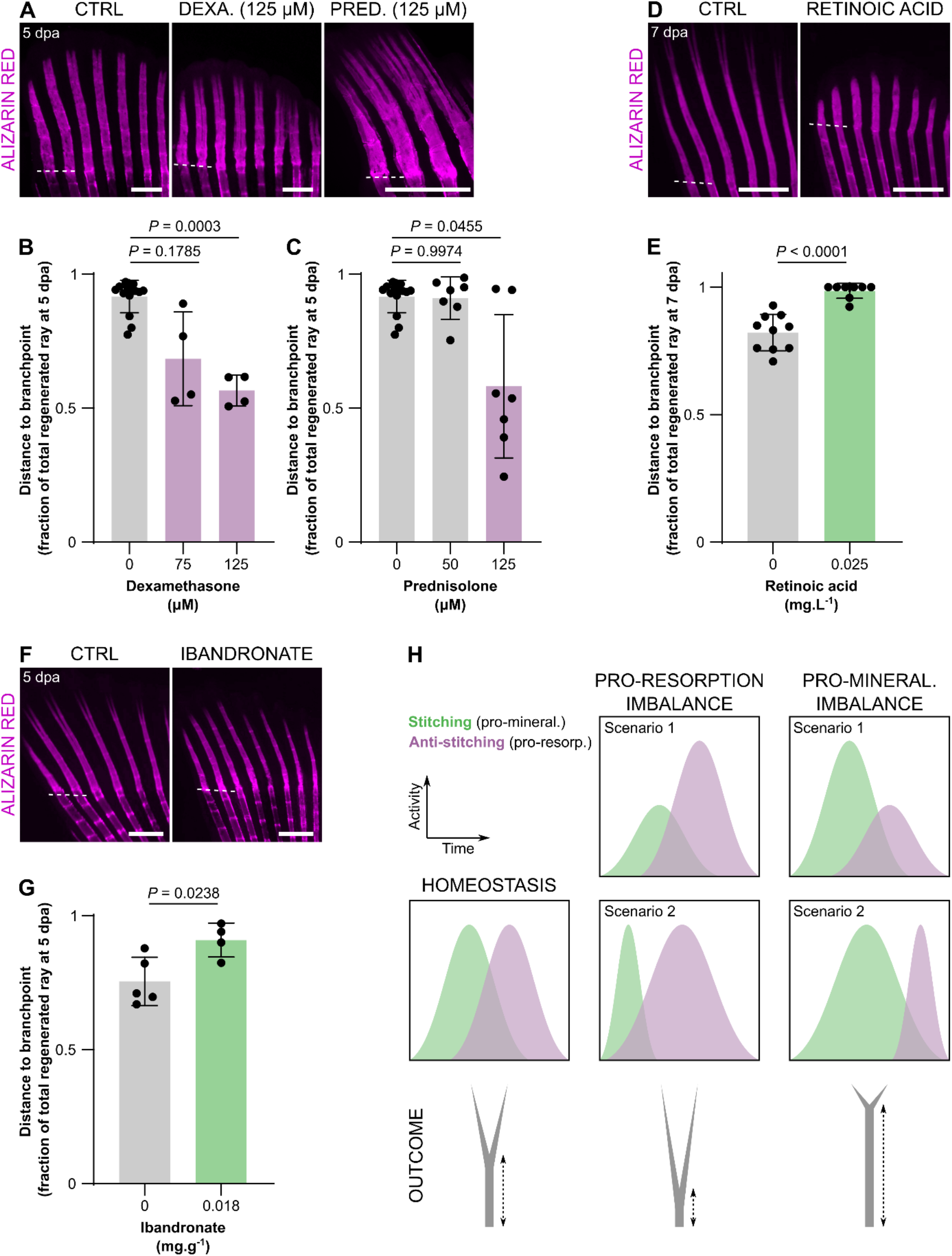
Branchpoint position, a proxy of ray bifurcation, is a fast readout of pro-resorption and pro-mineralization imbalances during bone formation. (**A-G**) Confocal images (A, D and F) and quantification (B, C, E, G) of mineralizing rays upon treatment with dexamethasone (A and B), prednisolone (A and C) retinoic acid (D and E) and ibandronate (F and G). (**H**) Model depicting how the position of the branchpoints represents pro-resorption and pro-mineralization imbalances. All graphs show the mean ± SD. Welch one-way ANOVA in B (*P* = 0.0004) and C (*P* = 0.0310) and Dunnett T3 post hoc tests for multiple comparisons; two-tailed Student’s t-test with Welch’s correction in E; two-tailed Student’s t-test in G. Scale bars: 500 μm.

Taken together, these data show that the branchpoint position reflects resorption and mineralization imbalances. Therefore, this model constitutes a fast readout to assess the properties of several agents during bone formation, considering the osteoblast-mediated stitching and OLT-mediated anti-stitching activities (Figure 6H).

## DISCUSSION

### Redefining the phases of ray mineralization in the regenerating zebrafish caudal fin

Despite the extensive knowledge on the mechanims regulating bone formation, much remains unknown on how bone size, shape, number and organization are defined. This process, known as bone patterning, has been widely studied in the vertebrate limb (Mariani and Martin, 2003). Although the teleost caudal fins have no homology with tetrapod appendages, they share many of the mechanisms activated during forelimb, hindlimb and digit development, including regulation by common signaling pathways such as Hox and Shh (Akimenko and Ekker, 1995; Nakamura et al., 2016; Yano and Tamura, 2013; Yano et al., 2012). Furthermore, the caudal fin regeneration model is a unique platform for live-imaging approaches to study the molecular and cellular events taking place during *de novo* bone formation in adult animals (Sehring and Weidinger, 2020).

So far, most studies have focused on the origins of osteoblasts, their dedifferentiation and redifferentiation, and spatial-temporal organization (Ando et al., 2017; Brandão et al., 2019; Knopf et al., 2011; Singh et al., 2012; Sousa et al., 2011; Watson and Kwon, 2015), without giving special attention to the mineralization process itself. Here, we focused on understanding how a single ray mineralizes and bifurcates. Shh has been shown to mediate the ray bifurcation process (Armstrong et al., 2017; Laforest et al., 1998; Quint et al., 2002; Zhang et al., 2012). It has been shown that ablation of *shha*-expressing epidermal cells leads to a delay in the branching of regenerating rays (Zhang et al., 2012). Moreover, ectopic expression of *shha* between the blastemas of two adjacent rays increases bone formation and ray fusion (Quint et al., 2002). More recently, *shha*-expressing cells were shown to escort pre-osteoblasts and position them to form the fin rays during regeneration.

Accordingly, the splitting of these domains induces the segregation of two equally discrete populations of pre-osteoblasts, thus driving ray branching (Armstrong et al., 2017). The regulatory role of Shh during ray bifurcation also involves the mediation of epithelial-mesenchymal interactions (Sehring and Weidinger, 2020). However, as previously suggested, Shh signaling may not be the sole responsible for ray bifurcation (Azevedo et al., 2012) thus raising the hypothesis that additional morphogenetic/cellular mechanisms are required. Using live-imaging approaches, we identified two mineralization fronts, indicating that the bifurcation process does not simply consist of the separation of one unit in two, but instead it results from the disruption of a fusion/stitching process between two pre-defined units.

Overall, we propose that ray bifurcation involves several stages. First, the morphogenetic action of molecules like Shh triggers the early division of the mineralizing ray by patterning pre-osteoblasts in two parallel domains on the flanks of the ray, as described by Armstrong et al. (2017). Second, the two mineralizing fronts fuse/stitch, through an osteoblast-dependent process. Third, an anti-stitching activity counteracts the fusion between the two mineralization fronts and defines the final position of the branchpoint.

Knowing that bone development and homeostasis result from the balanced activity of osteoblasts (i.e., to form new bone) and osteoclasts (i.e., to resorb bone), we hypothesized that osteoclasts or osteoclast-like cells could be actively mediating the anti-stitching process. Accordingly, we describe TRAP^+^ osteolytic tubules (OLTs) at the branching sites, that regulate the proximo-distal position of branchpoints. OLTs show a non-fluorescent lumen which suggests that they may be surrounding other structures, such as actinotrichia. In fact, actinotrichia have recently been sugested to have a role in ray bifurcation and in preventing ray fusion (Nakagawa et al., 2022). Another hypothesis is that OLTs are filled with a liquid substance, or possibly a reservoir of osteolytic enzymes, thus, explaining the accumulation of TRAP within the OLTs, as observed.

Osteoblasts and osteoclasts communicate through direct interaction or cytokine secretion to regulate cell differentiation, apoptosis and patterning (reviewed by Kim et al., 2020). Therefore, we hypothesize that osteoblasts recruit osteoclast-like cells to the branching sites through the secretion of molecules like M-CSF, RANKL/OPG, as it occurs during bone remodeling (reviewed by Kim et al., 2020). Osteoclasts are also known to secrete inhibitors of osteoblast differentiation, such as SEMA4D (Negishi-Koga et al., 2011), which would further enhance the anti-stitching properties of the OLTs.

The restoration of the original fin ray morphology during regeneration involves environmental cues from the surrounding tissue, in particular the interrays. For example, non-bifurcating peripheral rays are able to bifurcate when grafted between two central rays, and thus surrounded by two ectopic interrays (Murciano et al., 2002). On the other hand, swapped short and long rays regenerated into shorter or longer rays than the neighbouring host rays, indicating that there is an intrinsic local positional information driving the morphogenesis of these structures (Shibata et al., 2018). How OLTs are regulated by such environmental and local positional signals remains to be investigated.

Overall, we show that, in addition to the central role of osteoblasts in secreting and patterning the bone matrix, osteoclast-like cells are important players in defining the splitting of a single bony ray into two.

### The role of distinct osteoclast-like cells in multiple developmental and regenerative contexts

Osteoclasts are myeloid-derived cells (Burger, 1982; Walker, 1975) responsible for bone resorption through the acidification of the bone surface and the secretion of proteolytic enzymes (Henriksen et al., 2011; Teitelbaum, 2000, 2007; Väänänen and Zhao, 2008). These enzymes include cathepsin K (Ctsk; Brömme et al., 1996; Inaoka et al., 1995; Inui et al., 1997; Saftig et al., 1998), metalloproteinases (Kusano et al., 1998) and TRAP (Minkin, 1982; Price et al., 1995). TRAP is a *bona fide* marker used to assess osteoclast activity in mammals (Minkin, 1982; Price et al., 1995) and in teleosts, including zebrafish (Blum and Begemann, 2015b; Witten and Villwock, 1997).

Recent works brought a new perspective on osteoclasts and TRAP activity, which deviates from their classical role in bone resorption. As other monocytic cells, osteoclasts exhibit high plasticity depending on their environment and have the ability to modulate immune responses, including immunosuppression and inflammation (Madel et al., 2019). Osteoclasts may also play a non-bone-resorbing role during endochondral ossification, by controlling blood vessel anastomosis (Romeo et al., 2019), or in the development of bone lymphatics, possibly by carving paths for the lymphatic endothelial cells (Monroy et al., 2020). Furthermore, osteoclasts were shown to also express VEGF-C, a lymphatic growth factor (Zhang et al., 2008), thus they may regulate directly lymphatic formation within bone. In contrast, endothelial cells may replace osteoclast function in particular settings and secrete proteolytic enzymes essential for cartilage resorption and directional bone growth (Romeo et al., 2019). Moreover, lymphatic endothelial cells secrete colony-stimulating factor 1 (Csf1) to promote osteoclast formation and regulate bone resorption (Monroy et al., 2020). Non-canonical mechanisms of bone proteolysis, particularly those involving non-classical osteoclast-like cells and their role in regulating bone formation and shaping, are largely uncharacterized in zebrafish.

TRAP activity has been observed in the regenerative fin by Blum and Begemann (2015b). Yet, the authors used TRAP to infer on osteoclast activity but no molecular or cellular markers were used to identify the cells secreting this enzyme. Therefore, the precise localization of osteoclast-like cells, their morphology, their spatial-temporal dynamics, and their function during the regenerative process remained unknown. Here, we show that, during the pre-bifurcation phase, TRAP activity is associated with *ctsk*^+^ round osteoclast-like cells exhibiting cytoplasmic extensions, located mostly inside the regenerating rays. During the bifurcation phase, we show that TRAP-positive cells (i.e., OLTs), also positive for the *ctsk* reporter transgene, form tubular structures and align on the bone surface at the branching regions of the regenerating rays. Moreover, we found that OLTs are not specific to regeneration settings, but are more broadly associated with bone ray formation, such as during fin ray development.

Interestingly, these cells also express a reporter transgene for *lyve1b*, a widely used marker of lymphatic endothelial cells and fluorescent granular perithelial cells in zebrafish (e.g., Castranova et al., 2021; Okuda et al., 2012) that may also label some macrophages, as reported in mouse (Lim et al., 2018). Future work will address the exact origin of these cells and their association with the vascular and/or lymphatic systems. A recent work showed that lymphatics are required for proper cardiac regeneration in zebrafish (Gancz et al., 2019). The authors described isolated lymphatic sprouts in the injured area that are not connected with the lymphatic network of the uninjured tissue, suggesting that these cell clusters are specifically activated in response to injury. Therefore, it would be interesting to understand if a similar process takes place in the regenerating fin and how it relates with TRAP activity and bone shaping.

### Models to assess bone resorption-to-mineralization balance

Identification of compounds with osteogenic potential (i.e., pro-mineralogenic and/or anti-resorbing) can greatly contribute to the study of bone biology and to the establishment of new therapeutic targets to treat bone disorders or block disease progression.

Owing to the many advantages of using the zebrafish regenerating caudal fin model, (e.g., its suitability for the *in vivo* assessment of *de novo* bone formation using live-staining and live-imaging approaches (Bensimon-Brito et al., 2016), we and others have described technical strategies for drug screening with focus on bone mineralization (Cardeira et al., 2016; Laizé et al., 2014; Oppedal and Goldsmith, 2010; Recidoro et al., 2014). However, considering that a variety of bone diseases are the result of deregulated bone resorption, ranging from enhanced bone loss in osteoporosis to increased bone density in osteopetrosis (Bi et al., 2017), it is important to establish a straightforward model to study bone-resorbing activity and its balance with bone formation. Here we show that ray branchpoints are established during zebrafish caudal fin regeneration through a direct action of TRAP^+^ OLTs. We also show that inhibition of bone-resorbing activity is sufficient to distalize the branchpoints, which can be used as clear readouts of resorption-to-mineralization imbalances (Fig. 6). Accordingly, several reports have documented alterations branchpoints positions in the zebrafish regenerating caudal fins rays. A retinoic acid-induced inhibition of bifurcation was previously proposed by White et al. (1994), an effect that we reproduced and show in detail in the present study. Branchpoints were distalized upon inhibition of the Calcineurin pathway (Kujawski et al., 2014), known to promote osteoclastogenesis (Sun et al., 2007). Recently, we showed that an anti-mineralization/pro-resorption imbalance induced by benzo[α]pyrene resulted in proximalized branchpoints (Tarasco et al., 2021). Ablation of *mpeg1*-positive macrophages, a potential source and/or regulator of the OLTs, resulted in a reduced number of bifurcated rays (Petrie et al., 2014). Interestingly, a recent study proposed that ray branching is modulated by biomechanical forces (Dagenais et al., 2021). In fact, bone formation is well known to rely on the mechanosensitive properties of bone-forming cells that, in addition to bone deposition, are also involved in the recruitment of osteoclasts and, therefore, regulate bone resorption (Xiao and Quarles, 2015).

Overall, we propose that branchpoint positioning could serve as a simple and straightforward system to study the (im)balance between bone resorption and mineralization activities. Ultimately, this system will provide a unique platform to study different compounds, signaling pathways and molecular mechanisms, towards the identification of new targets and the development of therapeutic drug strategies.

## METHODS

### Animals

All experiments were performed using adult zebrafish (3-6 months old) or juvenile fish (up to 3 weeks old). Animal husbandry followed standard conditions. All animal procedures followed institutional (CCMAR) guidelines, and the European and Portuguese legislation for animal experimentation and welfare (Directives 86/609/CEE and 2010/63/EU; Portaria 1005/92, 466/95 and 1131/97; Decreto-Lei 113/2013). Animal handling and experimentation were performed by qualified operators accredited by the Portuguese Direção-Geral de Alimentação e Veterinária (authorization no. 012769). All efforts were made to minimize pain, distress, and discomfort. Anesthesia was performed by incubating fish in 0.6 mM tricaine solution (MS-222; Sigma-Aldrich, St. Louis, MO, USA). Euthanasia was performed using a lethal dose of anesthetic.

Wild-type and transgenic zebrafish used in this study were from the AB strain. The lines used in this study were: *Tg(Ola*.*bglap:EGFP)*^*hu4008*^ (Knopf et al., 2011), abbreviated *bglap:GFP*; *Tg(sp7:mCherry-Eco*.*NfsB)*^*pd46*^ (Singh et al., 2012), abbreviated *sp7:mCherry-NTR*; *Tg(Ola*.*ctsk:FRT-DsRed-FRT-Cre,myl7:EGFP)*^*mh201*^ (Caetano-Lopes et al., 2020), abbreviated *ctsk:DsRed*; *Tg(mpeg1*.*1:NTR-EYFP)*^*w202*^ (Petrie et al., 2014), abbreviated *mpeg1*.*1:YFP*; *Tg(fli1:EGFP)*^*y1*^ (Lawson and Weinstein, 2002), abbreviated *fli1:GFP*; *Tg(−5*.*2lyve1b:EGFP)*^*nz150*^ (Okuda et al., 2012), abbreviated *lyve1b:GFP*; *Tg(−2*.*4shha-ABC:GFP)*^*sb15*^ (Shkumatava et al., 2004), abbreviated *shha:GFP*.

### Caudal fin amputation

Amputation was performed 1-2 segments below the branchpoint of the most peripheral branching rays in anesthetized fish. Upon amputation, fish were transferred to isolated containers at a maximum density of 5.5 fish/l. Water and light conditions were adjusted to match those of the rearing system, according to the standard conditions of zebrafish husbandry. All fin regeneration experiments took place at 33ºC, a standard temperature in regeneration studies (Cardeira et al., 2016; Nechiporuk and Keating, 2002; Oppedal and Goldsmith, 2010).

### Transcriptome analysis

Regenerated tissue was homogenized in solution D (4 M guanidinium thiocyanate, 25 mM sodium citrate pH 7.0, 0.5% N-laurosylsarcosine, 0.1 M 2-mercaptoethanol; all in DEPC-treated water) using a 20 G needle and total RNA was extracted through phenolic extraction as previously described (Chomczynski and Sacchi, 2006). Briefly, 1 volume of phenol at pH 4.5 and 0.2 volume of chloroform/isoamyl alcohol mixture (49:1) was added to the homogenate (all chemicals from Sigma-Aldrich), and the mixture was inverted 2-3 times and cooled for 10 min on ice. The mixture was centrifuged at 10000 *g* for 15 min and the aqueous phase transferred to a new tube. RNA solution was further purified using 1 volume of chloroform/isoamyl alcohol mixture, as described above. Total RNA was precipitated by adding 1 volume of ice-cold 2-propanol (Sigma-Aldrich) to the aqueous phase and incubating the mixture for 24 h at −80ºC. Precipitated RNA was pelleted (centrifugation at 12000 *g* for 10 min) and the supernatant removed. The RNA pellet was washed once with 75% ethanol and further centrifuged at 14000 *g* for 10 min. Ethanol was removed and the RNA pellet was air-dried at room temperature and resuspended in 30 μL of RNAse-free water (Sigma-Aldrich). All centrifugations were performed at 4ºC. Quantity and quality of the RNA were assessed using an Experion electrophoresis system (Bio-Rad, Hercules, CA, USA) and samples with an RNA integrity number (RIN) > 8 were further processed for RNA-Seq.

The construction of cDNA libraries was carried out using the Illumina TruSeq Stranded mRNA Library Preparation kit. cDNA fragments were sequenced using the Illumina Hiseq 2500 platform with 100 bp paired-end sequencing reads.

Raw sequences were trimmed to generate high quality data using the CLC Genomics Workbench 9.0.1, as follows: quality trimming based on quality scores (0.01), ambiguity trimming (2 nucleotides) and length trimming (minimum of 30bp). After quality and length trimming, base trim – to remove a specified number of bases at either 3’ or 5’ ends of the reads – was found unnecessary. For each original read, the regions of the sequence to be removed were determined independently of each type of trimming operation. Mapping of the reads was performed against the *Danio rerio* reference genome (assembly GRCz10) with length (minimum percentage of the total alignment length that must match the reference sequence at the selected similarity fraction) and similarity (minimum percentage identity between the aligned region of the read and the reference sequence) parameters set to 0.95. Gene expression was calculated based on the Reads per Kilobase of exon model per Million mapped reads (RPKM) approach (Mortazavi et al., 2008).

Expression levels were calculated using the RPKM values from each sample independently. Differential expression was then calculated using a multi-factorial statistical analysis based on a negative binomial model that used a generalized linear model approach influenced by the multi-factorial EdgeR method (Robinson et al., 2010). The differentially expressed genes were filtered using standard conditions (Raza and Mishra, 2012; Robinson et al., 2010), i.e., a False Discovery Rate (FDR) *P* value <0.05 and a fold change >2 or <-2.

The quality of the produced data was ensured by evaluating the Phred quality score at each cycle (position in read; ensuring a minimum Phred score of 20). Further quality control was performed by principal component analysis (PCA), hierarchical clustering (considering Euclidean distance) and heat map analysis.

Gene ontology analysis was performed using PANTHER (Protein ANalysis THrough Evolutionary Relationships; Mi et al., 2016) v16.0. For this analysis, Ensembl IDs of the genes upregulated at 3 dpa were used as input. Fisher’s Exact test was performed to analyze for overrepresentation, using the Reactome pathway annotation set and the *Danio rerio* genome as a reference list. For the generation of the gene expression heat maps, Heatmapper (www.heatmapper.ca; University of Alberta) was used. Differential expression of selected genes was transformed to a Z-score per row and clustered using the average linkage clustering and the Euclidean distance measuring methods. The list of collagens, ECM regulators and other structural ECM components was retrieved from the zebrafish *in silico* matrisome (Nauroy et al., 2018).

### Staining and immunohistochemistry

Mineral staining was performed in live animals following established protocols using alizarin red S (ARS; Bensimon-Brito et al., 2016) or calcein (Du et al., 2001), according to the need for combination with red or green fluorophores. Briefly, animals were incubated in system water containing 0.01% ARS or 0.2% calcein for up to 15 min, and rinsed at least 3 times in clean system water. Imaging was performed in fins immediately after collection from euthanized fish, or in anesthetized animals, depending on the need to track mineralization over time.

To assess bone-resorbing activity, the activity of tartrate resistant acid phosphatase (TRAP) was determined, as described (Blum and Begemann, 2015b). Briefly, fins were fixed in 4% PFA for 3 hours at room temperature, washed 5 times in PBT for 5 min and incubated in PBTx for 30 min. The samples were then incubated in TRAP buffer (50mM sodium tartrate, 0.1 M acetic acid and 0.1 M sodium acetate; pH 4.4; all reagents were from Sigma-Aldrich) for approximately 20 min. The colorimetric assay was performed by incubating the samples in TRAP buffer containing 0.1 mg/ml Naphtol AS-MX phosphate and 0.3 mg/ml Fast Red Violet LB. The samples were washed in 1× PBS for 5 min and cleared with 1.5% KOH for 5 min. The samples were then gradually transferred to 70% glycerol in 1× PBS and imaged under bright field or fluorescence microscopy.

Immunohistochemistry was performed immediately after staining for TRAP activity before transferring to glycerol. For that purpose, DsRed was immunostained in *ctsk:DsRed* fish using a DsRed polyclonal primary antibody raised in rabbit (Living Colors DsRed, Cat. No. 632496; Takara, Bio Inc., Kusatsu, Shiga, Japan) and anti-rabbit Alexa Fluor 488 secondary antibody raised in goat (Invitrogen, Waltham, MA, USA).

### Ablation and drug treatments

For osteoblast ablation assays, *sp7:*mCherry-NTR transgenic zebrafish were let to regenerate until 3 dpa. Fish were then incubated in a Metronidazole (Mtz) solution for 24 hours. Mtz (Sigma-Aldrich) was freshly prepared in dimethylsulfoxid (DMSO, Sigma-Aldrich) and diluted in system water to a final concentration of 8.5mM with 0.2% of DMSO. Control fish were incubated in system water with 0.2 % DMSO alone. Both Mtz and vehicle-treated fish were maintained for 24 h in the dark to prevent Mtz degradation (Curado et al., 2008). Fish were then rinsed twice to washout any traces of Mtz and returned to system water until 7 dpa. Mtz and vehicle-treated fish were anaesthetized and regenerates were collected and labelled with calcein to visualize the calcified bony-rays.

Salmon calcitonin (Sigma-Aldrich) was prepared in 1× PBS and administered at 0.5, 5 and 50 μg g^-1^ through a single intraperitoneal injection at the time of fin amputation. Briefly, fish were dried on absorbent paper and weighed immediately before the injection. A volume of 30 μl per gram of fish was injected; injection solutions were adjusted to the desired concentrations accordingly. Control fish were injected with the vehicle (1× PBS).

For the other drugs, fish were continuously exposed by immersion from the moment of fin amputation until the experimental endpoints. Dexamethasone (75 and 125 μM; Sigma-Aldrich), prednisolone (50 and 125 μM; Sigma-Aldrich) and all-trans retinoic acid (0.025 mg l^-1^; Sigma-Aldrich) were prepared in to achieve a working concentration of 0.1% DMSO; control fish were exposed to 0.1% DMSO alone. Ibandronate (0.018 mg g^-1^; Roche, Basel, Switzerland) was prepared in system water; control fish were placed in system water alone. 70% of the treatments were renewed daily.

### Microscopy and imaging

As mentioned above, imaging of adult fish was performed under anesthesia or on collected fins, following animal euthanasia. For juvenile imaging, anesthetized fish were placed directly under a Zeiss LSM800 Observer (Zeiss) inverted confocal microscope and ZEN 3.1 (Blue edition) software. For general histomorphometry, ARS stained fins were imaged under fluorescence conditions using a Leica MZ 7.5 fluorescence stereomicroscope (Leica Microsystems GmbH, Wetzlar, Germany), coupled to an F-View II camera controlled by the Cell^F v2.7 software (Olympus Soft Imaging Solutions GmbH, Münster, Germany), or using an MZ10F fluorescence stereomicroscope (Leica Microsystems GmbH), equipped with a Leica DFC7000T color camera. For bright field imaging, a SteREO Lumar.V12 (Zeiss, Oberkochen, Germany) or a Nikon SMZ25 coupled with Nikon Digital Sight DS-Ri1 camera (Nikon, Minato City, Tokyo, Japan), were used. For detailed analyses, samples were imaged using a Zeiss LSM700 (Zeiss) or LSM800 inverted confocal microscope. For the osteoblast ablation studies, fins were imaged under a Zeiss Axio Observer z1 inverted microscope equipped with an axiocam 506 monochromatic camera, using an EC Plan-Neofluar 5x 0.16NA air objective controlled by Zen 3 blue software.

Z projections of maximum intensity and orthogonal projections were performed using ImageJ v1.51n (Wayne Rasband, National Institutes of Health, Bethesda, MD, USA) or ZEN 3.1 (Blue edition). 3D reconstructions were performed using the Imaris (Bitplane) software.

### Morphometry, quantitative analyses and statistics

The distance to branchpoint was determined by calculating the fraction of the length from the amputation plane to the branchpoint relative to the total mineralized length (from the amputation plane to the distal-most mineralized point). The mean of the relative distances to branchpoint of rays #2, 3, 4 and 5 (peripheral-most bifurcating rays; Cardeira et al., 2016)) from each lobe of each fish were used in all comparative analyses. For the mineralization and branchpoint tracking analysis in single rays, ray #3 from the dorsal lobe of each fish was measured. For quantification of TRAP activity, the number of rays displaying TRAP signal at the distal tip, coinciding with the branching area, was counted and calculated as percentage of the total number of rays within each fin. *shh:*GFP^+^ domains were evaluated by measuring their proximo-distal length divided by the total mineralized ray length (from the amputation plane to the distal-most mineralized point).

Statistical analyses were performed using Prism v8.4.3 (GraphPad Software Inc., La Jolla, CA, USA). Two-group comparisons were conducted using the unpaired, two-tailed Student’s t-test when groups followed a normal distribution. Unpaired, two-tailed Student’s t-test with Welch’s correction was applied for comparisons using samples without normal distribution. Multiple comparisons were conducted using one-way ANOVA and Tukey’s post-hoc tests. For comparisons of repeated measures, one-way ANOVA with Geisser-Greenhouse correction and Tukey’s post-hoc tests were used. For comparisons with samples with significantly different standard deviations, Welch one-way ANOVA and Dunnett T3 post hoc tests were used. Normality was tested through the Shapiro-Wilk’s test. Bartlett’s test was used to determine if samples display equal variances in multiple comparisons. Significance level was set to α=0.05 for all tests.

## ACKNOWLEDGEMENTS

This study was funded by the Portuguese Foundation for Science and Technology (FCT) through the project UIDB/04326/2020 (to the CCMAR) and grant PD/BD/52425/2013 (to J.C.-S.). Research in the Stainier lab is funded in part by the Max Planck Society. We thank Matthew Harris and Joana Caetano-Lopes for kindly providing the *ctsk*:*DsRed* transgenic line.

## AUTHOR CONTRIBUTIONS

Conceptualization and experimental design, J.C.-S., A.B.-B., P.J.G., D.Y.R.S. and V.L.; Experimental Execution, J.C.-S., A.B.-B., M.T., A.S.B., J.R. and P.J.A.; RNA-Seq analysis, J.C.-S. and P.J.A.; Morphometric analysis, J.C.-S.; Imaging analysis, J.C.-S., A.B.-B., M.T., A.S.B. and J.R; Drug experiments, J.C.-S., A.B.-B., M.T., J.R., A.S.B.; Resources, A.J., M.L.C., P.J.G., D.Y.R.S. and V.L.; Writing – Original Draft, J.C.-S. and A.B.-B.; Writing – Review & Editing, all authors; Project Administration, J.C.-S., A.B.-B., P.J.G., D.Y.R.S. and V.L.; Funding Acquisition, P.J.G., D.Y.R.S. and V.L.

## DECLARATION OF INTERESTS

The authors declare no competing interests.

## Notes

### Competing Interest Statement

The authors have declared no competing interest.

