## Supplementary figure 1 for "Fin ray branching is defined by TRAP^+^ osteolytic tubules"

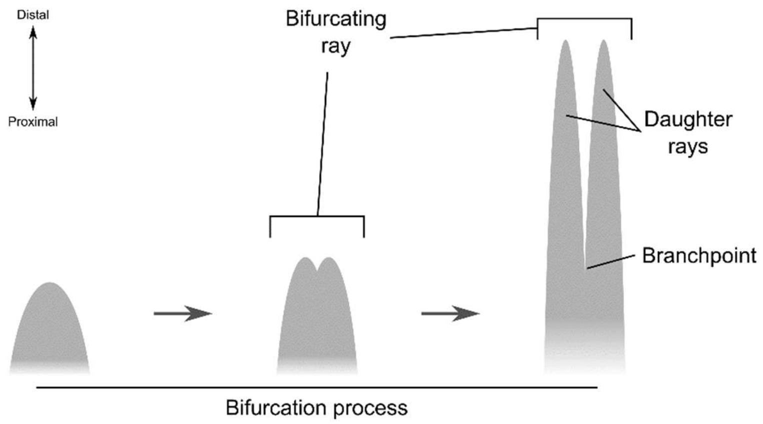


**Figure S1. Simple scheme of the ray bifurcation process.** The gray shapes represent the mineralized regenerating bony rays. A bifurcating ray display two visible daughter rays forming distally. The branchpoint is the proximal-most point of daughter ray splitting.
