## Supplementary figure 2 for "Fin ray branching is defined by TRAP^+^ osteolytic tubules"

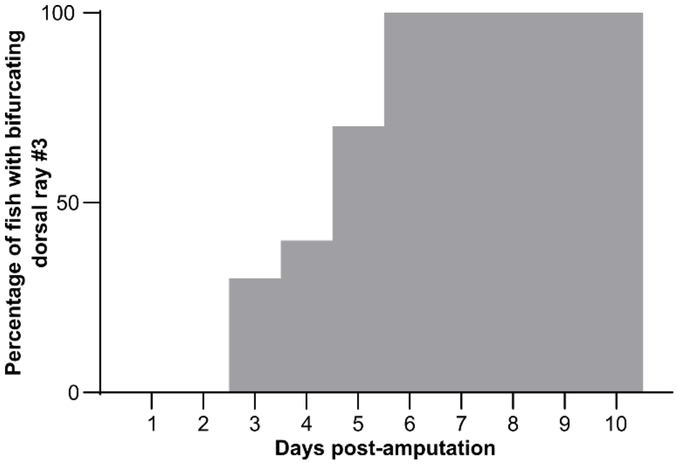


**Figure S2. Percentage of fish exhibiting bifurcating ray.** The dorsal ray #3 of each fish was used for the analysis (*N* = 10). Based on the same data as for Figure 1B.
