## Supplementary figure 3 for "Fin ray branching is defined by TRAP^+^ osteolytic tubules"

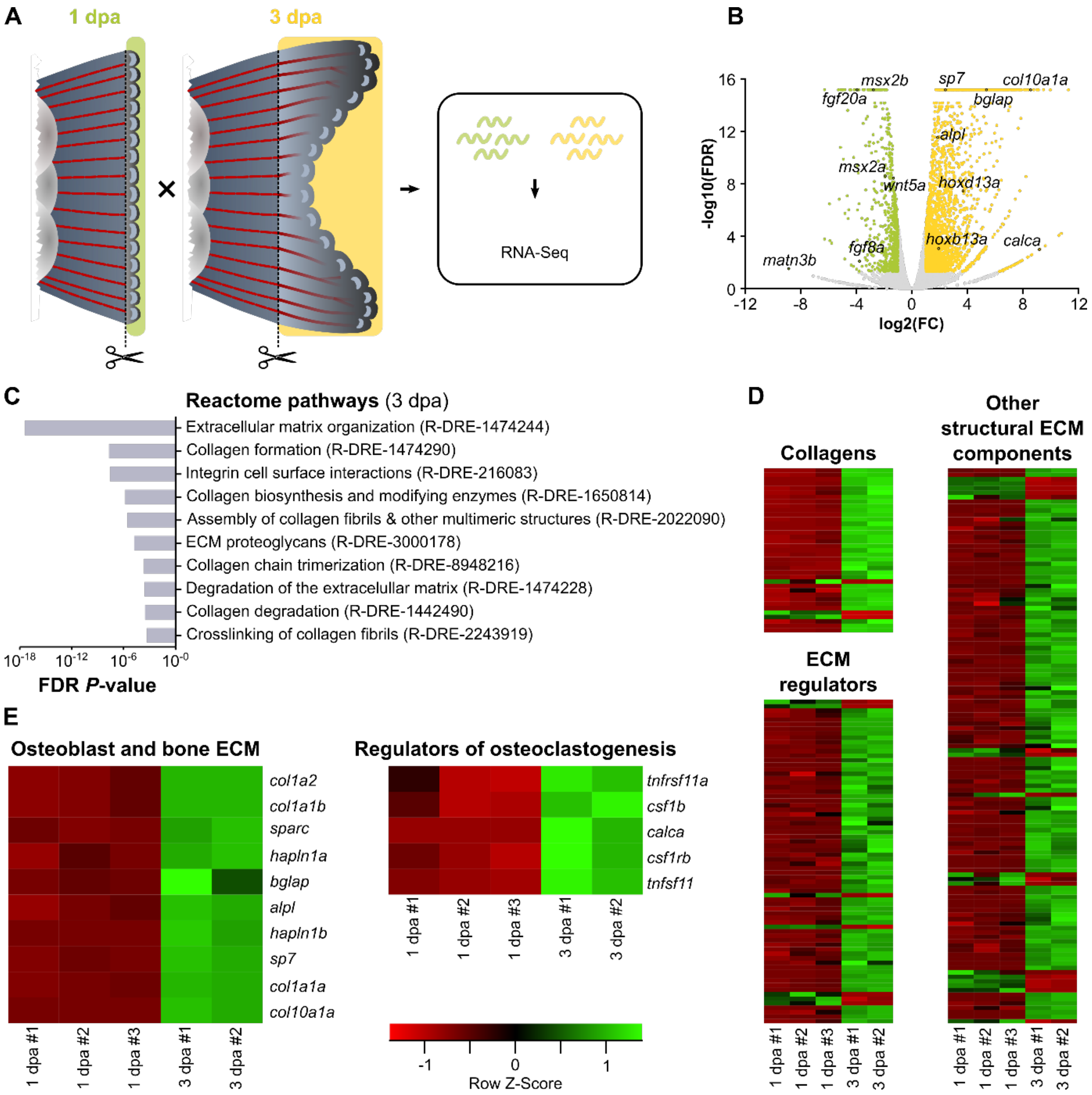


**Figure S3. RNA-Seq analysis reveals induction of osteoblast- and osteoclast-related genes, and activation of morphogenetic and ECM remodeling processes in regenerates preceding bifurcation**. (**A**) Experimental design for RNA-Seq. (**B**) Volcano plot showing differentially expressed genes (FC ≥ 2; FDR < 0.05). Green and yellow represent genes with enriched expression at 1 and 3 dpa, respectively. (**C**) Top 10 overrepresented gene ontology terms, represented by Reactome pathways, among the list of upregulated genes at 3 dpa. The Reactome identifiers are depicted. (**D, E**) Heatmaps for individual samples, showing the relative expression of genes encoding collagens, ECM regulators and other structural ECM components (**D**), based on the zebrafish matrisome (Nauroy et al., 2018) and of osteoblast-specific genes, bone ECM genes and genes involved in osteoclastogenesis (**E**).
