## Supplementary figure 4 for "Fin ray branching is defined by TRAP^+^ osteolytic tubules"

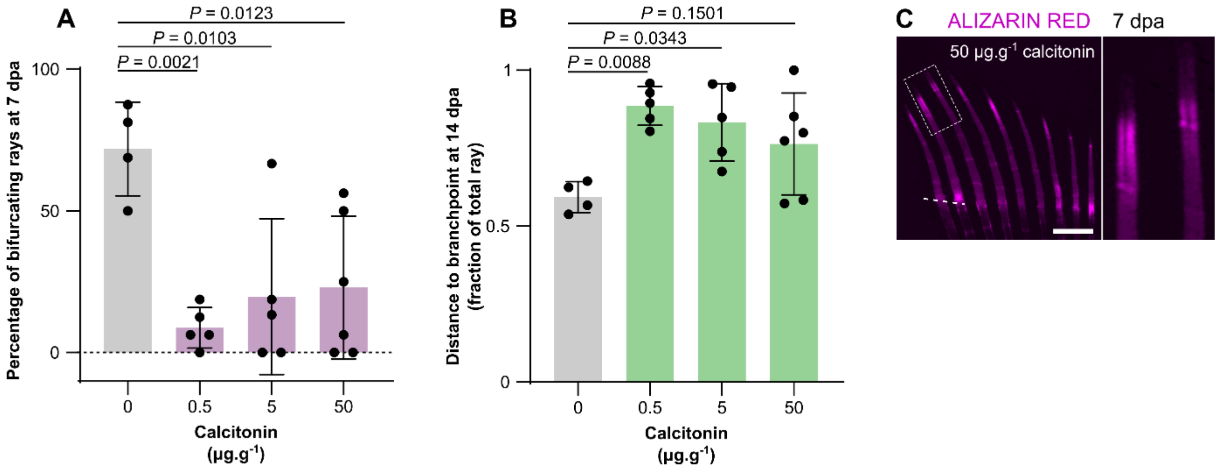


**Figure S4. Additional analyses further strengthen the bifurcation phenotype upon osteoclast activity inhibition.** (**A**) Quantification of the percentage of bifurcating rays at 7 dpa. (**B**) Relative distance from the amputation plane to the branchpoint at 14 dpa. (**C**) Example stereomicroscope image of mineralized rays from a fin at a bifurcation stage of fish treated with the highest calcitonin concentration (50 µg.g$^{-1}$), showing impaired bifurcation and heterogeneities in mineralization. All graphs show the mean ± SD. One-way ANOVA ($P$ = 0.0025 in A and $P$ = 0.0111 in B) and Tukey's post-hoc test. Scale bar: 500 µm.
