## Supplementary table 1 for "Fin ray branching is defined by TRAP^+^ osteolytic tubules"

**Table S1.** Statistics details on Tukey's multiple comparisons test related to Figure 1B.

| Tukey's multiple comparisons test | Summary | Adjusted *P* Value |
|---|---|---|
| 3 dpa vs. 4 dpa | Non-significant | 0.4099 |
| 3 dpa vs. 5 dpa | Non-significant | 0.1575 |
| 3 dpa vs. 6 dpa | Non-significant | 0.0782 |
| 3 dpa vs. 7 dpa | * | 0.0461 |
| 3 dpa vs. 8 dpa | * | 0.0474 |
| 3 dpa vs. 9 dpa | * | 0.0247 |
| 3 dpa vs. 10 dpa | Non-significant | 0.0512 |
| 4 dpa vs. 5 dpa | Non-significant | 0.2894 |
| 4 dpa vs. 6 dpa | ** | 0.0072 |
| 4 dpa vs. 7 dpa | * | 0.0254 |
| 4 dpa vs. 8 dpa | ** | 0.0033 |
| 4 dpa vs. 9 dpa | ** | 0.0066 |
| 4 dpa vs. 10 dpa | ** | 0.0014 |
| 5 dpa vs. 6 dpa | Non-significant | 0.1323 |
| 5 dpa vs. 7 dpa | * | 0.0285 |
| 5 dpa vs. 8 dpa | * | 0.0182 |
| 5 dpa vs. 9 dpa | ** | 0.0064 |
| 5 dpa vs. 10 dpa | ** | 0.0025 |
| 6 dpa vs. 7 dpa | Non-significant | 0.9758 |
| 6 dpa vs. 8 dpa | Non-significant | 0.1658 |
| 6 dpa vs. 9 dpa | Non-significant | 0.2537 |
| 6 dpa vs. 10 dpa | Non-significant | 0.0508 |
| 7 dpa vs. 8 dpa | Non-significant | 0.4118 |
| 7 dpa vs. 9 dpa | Non-significant | 0.2558 |
| 7 dpa vs. 10 dpa | Non-significant | 0.0984 |
| 8 dpa vs. 9 dpa | Non-significant | >0,9999 |
| 8 dpa vs. 10 dpa | Non-significant | 0.3668 |
| 9 dpa vs. 10 dpa | Non-significant | 0.4733 |
